# *Kabasura Kudineer Choornam*, a medicinal polyherbal formulation, modulates human macrophage polarization and phagocytic function

**DOI:** 10.1101/2024.12.20.629574

**Authors:** Asli Korkmaz, Duygu Unuvar, Sinem Gunalp, Derya Goksu Helvaci, Duygu Sag

## Abstract

**Background:** *Kabasura kudineer choornam* (*KKC*) is a polyherbal formulation with 15 ingredients. It has been shown to have anti-inflammatory and anti-microbial properties and demonstrate efficacy in managing the symptoms of H1N1 swine flu and COVID-19. However, its mechanism of action is not fully comprehended.

**Purpose:** Herein, we examined the effect of the *KKC* on the polarization and function of primary human macrophages.

**Methods:** Human monocyte-derived macrophages (M0 macrophages) pre-treated with *KKC* extract were polarized into M1, M2a, or M2c subtypes. The expression of the M1/M2 polarization markers was analyzed by qPCR, flow cytometry, and ELISA, and the phagocytosis capacity of macrophages was analyzed by flow cytometry.

**Results:** Our data show that the *KKC* treatment increased the expression of the M1 markers IDO1, IL-1β, IL-12a(p35), and TNF in both polarized and unpolarized macrophages at mRNA level. However, it decreased the secretion of IL-12 (p70) in M1 macrophages and increased the secretion of TNF in M0, M2a, and M2c macrophages. IL-10 secretion was increased in M0 and M2a macrophages, while it was decreased in M1 macrophages after the *KKC* treatment. Interestingly, all *KKC*-treated macrophage phenotypes displayed a downregulation in the expression of the M1/M2 surface markers CD64, CD206, CD209, and CD163, which also play a role in phagocytosis. In accordance with this result, the phagocytic capacity of both polarized and unpolarized macrophages was decreased after the *KKC* treatment.

**Conclusion:** In conclusion, *KKC* extract modulates macrophage inflammatory response and could be a potential supplement for the treatment of infectious and inflammatory diseases.

## 1. Introduction

Studies have demonstrated the preventive and therapeutic benefits of plants and their compounds in addressing a spectrum of common and complex diseases including cancer and various infectious diseases (Dhama et al., 2015, 2014, 2014, 2013; Mahesh et al., 2015; Mahima et al., 2012; Rahal et al., 2014; Singh et al., 2016). The *Kabasura kudineer choornam* (*KKC*) is a polyherbal formulation derived from Siddha medicine originating in India (Parameswaran et al., 2021). The *KKC* formulation contains 15 different components (Behera et al., 2023; John et al., 2015; Mekala and Murthy, 2020) (Table1) and is commonly recommended for the efficient management of prevalent respiratory conditions including colds, coughs, respiratory distress, and influenza-like symptoms (John et al., 2015; Mekala and Murthy, 2020). The *KKC* has been reported to have anti-inflammatory, anti-viral, anti-pyretic, immunomodulatory and anti-bacterial properties due to its phytochemical components (Parameswaran et al., 2021; Saravanan et al., 2018; Thillaivanan et al., 2015). It has been prescribed against H1N1 swine flu and has demonstrated great efficacy in managing the disease symptoms and enhancing the body’s immunological response (Thillaivanan et al., 2015).

**Table 1.** The components of *Kabasura kudineer choornam*.

| Cp. # | Botanical Name | Local Name | Common Name | Part |
| --- | --- | --- | --- | --- |
| 1 | *Zingiber officinale* Roscoe | Chukku | Ginger | Rhizome |
| 2 | *Piper longum* L. | Thippili / Pippali | Long pepper | Fruit |
| 3 | *Syzygium aromaticum* (L.) Merr. & L.M.Perry | Kirambu | Clove | Flower bud |
| 4 | *Tragia involucrate* L. | Sirukanchori | Indian stinging nettle, climbing nettle | Root |
| 5 | *Anacyclus pyrethrum* (L.) Lag. | Akkirakaram | Pellitory | Root |
| 6 | *Hygrophila auriculata* (Schumach.) Heine | Mulliver | Vajradanti | Root |
| 7 | *Terminalia chebula* var. | Kadukkaithol | Yellow myroblan | Pericarp |
| 8 | *Adhatoda vasica* L. | Adathodai / Vasaka | Adathodai | Leaf |
| 9 | *Coleus amboinicus* Lour | Karpuravalli | Bishop's weed | Leaf |
| 10 | *Saussurea lappa* (Decne.) Sch. Bip. | Kostam | Spiral Ginger | Root |
| 11 | *Tinospora cordifolia* (Willd.) Miers | Guduchi | Heart-leaved moonseed | Stem |
| 12 | *Clerodendron serratum* (L.) Moon | Siruthekku / Bhrangi | Beetle killer | Root |
| 13 | *Andrographis paniculata* (Burm.f.) Nees | Nilavmebu / Bhunimba | Andrographis | Whole plant |
| 14 | *Sida acuta* Burm.f. | Vattatthiruppi | Morning mallow | Root |
| 15 | *Cyperus rotundus* L. | Korai kizhangu | Cocgrass | Rhizome |

After the recent COVID-19 outbreak, in-silico screening methods and molecular docking studies indicate that the parent compounds from the *KKC* may exhibit anti-viral effects against SARS-CoV-2 by blocking the host cell receptor -ACE2 or inhibiting the key viral protease necessary for its replication in the host cell (Kiran et al., 2020). This action could potentially inhibit COVID-19 more effectively than synthetic drugs due to a superior energy score (Kiran et al., 2020; Nallusamy et al., 2021; Vincent et al., 2020). Different randomized clinical trials on COVID-19 patients have shown that *Kabasura kudineer* as an adjunct treatment, reduced SARS-CoV-2 viral burden, prevented progression from asymptomatic to symptomatic state, improved symptoms, led to changes in immune markers and decreased hospitalization length in COVID-19 patients without serious adverse events and mortality (Chitra et al., 2021; Jamuna et al., 2020; Meenakumari et al., 2021; Natarajan et al., 2021; Rao et al., 2020; Srivastava et al., 2021). As a result, the *KKC* is recommended as a therapeutic medicine for enhancing the immune system and combating COVID-19, as per the Ministry of AYUSH guidelines issued by the Government of India (AYUSH Ministry of Health Corona Advisory—D.O. No. S. 16030/18/2019—NAM; 06th March 2020) (Jose et al., 2022).

Macrophages are innate immune cells of the myeloid lineage. They originate from blood monocytes, which differentiate into either pro-inflammatory M1 (classically activated) or anti-inflammatory M2 (alternatively activated) macrophages in presence of certain polarization factors when recruited into peripheral tissue (Davis et al., 2013; Gordon and Taylor, 2005; Italiani and Boraschi, 2014; Yang et al., 2014). Macrophages can be polarized into M1 phenotype by IFN-γ/LPS or TNF *in vitro* (Mantovani et al., 2005). M2 macrophages are categorized into three subgroups: M2a, M2b, and M2c, depending on the stimuli they respond to and the markers they express (Murray et al., 2014). M2a polarization is induced by IL-4 and/or IL13; M2b polarization is induced by immune complex and FcϒR/TLR ligands; and M2c polarization is induced by IL-10, TGFβ or glucocorticoid stimulation (Gordon, 2003). M1 macrophages support the anti-microbial and anti-tumoral response, while M2 macrophages are involved in tissue healing, allergic reactions or cell proliferation according to their subgroup (Mantovani et al., 2004; Munder et al., 1998). Imbalance between M1/M2 macrophage phenotypes underlies numerous diseases. For instance, M1 macrophages in type 2 diabetes and atherosclerosis promote inflammation, obesity, and insulin resistance, while M2-like macrophages alleviate these symptoms in symptomatic plaques (Espinoza-Jiménez et al., 2012; Gaetano et al., 2016; Lumeng et al., 2007; Menghini et al., 2012). Moreover, M1 macrophages have been linked to a strong type I interferon response and elevated disease severity and mortality in infections caused by viruses like SARS-CoV-2 (Feng et al., 2020; Lv et al., 2021; Mehta et al., 2020; Merad and Martin, 2020; Ziegler et al., 2020) and the highly virulent H5N1 influenza A virus (Baskin et al., 2009; Channappanavar et al., 2016).

Although *KKC* is known to enhance the body’s immune response, its mechanism of action is not well understood. In this study, we investigated the impact of *KKC* on polarization and function of human macrophages. Our results demonstrate that *KKC* modulates polarization and decreases phagocytic function of primary human macrophages with a downregulation of certain phagocytic surface receptors.

## 2. Materials and Methods

### 2.1. Study approval

Buffy coats from healthy donors were obtained from Dokuz Eylul University Hospital Blood bank (Izmir, Turkey) after written consent. Ethical approval was granted by the Non-Interventional Research Ethics Committee of Izmir Biomedicine and Genome Center (Izmir, Turkey) for the use of buffy coats (Approval number: 2021-042).

### 2.2. Primary human monocyte isolation and macrophage differentiation

Primary human monocytes were isolated from buffy coats by gradient centrifugation technique, using Ficoll-Paque (GE healthcare) followed by Percoll (GE healthcare) (Menck et al., 2014). Isolated primary monocytes were then cultured in RPMI-1640 medium (Gibco) completed with 5% heat-inactivated fetal bovine serum (FBS) (Gibco) and 1% Penicillin-Streptomycin (Gibco) (R5 medium), supplemented with 10 ng/mL recombinant human M-CSF (PeproTech) in ultra-low attachment six-well plates (Corning) for 7 days at 37°C and 5% CO_2_ (Gunalp et al., 2023). After 7 days, the cells were collected, and the macrophages were verified to be approximately 90% CD68+ by flow cytometry.

### 2.3. Extraction and preparation of KKC

The *KKC* (powder form) was kindly provided by Dr. Ravi Kumar Reddy from Sri Sri Tattva, (Bengaluru, Karnataka, India). As *KKC* contains active compounds including alkaloids, carbohydrates, glycosides, flavonoids, phenols, saponins, tannins, and terpenoids and also in small quantities of flobatannins, essential oils, vitamin C, proteins, and amino acids derived from diverse medicinal plants (Mekala and Murthy, 2020), we obtained *KKC* extract by dissolving the fine powder in DMSO, due to its ability to dissolve both polar and non-polar compounds (Kuroda et al., 2020). Stock solution concentration was set to 100 mg/ml. Following 6 hours of gentle agitation and 18 hours of upright settling at room temperature (RT), the solution was centrifuged at 200 g for 5 minutes. The supernatants were kept at -80°C for future use.

### 2.4. Treatments

Human monocyte derived macrophages were plated into 24-well plates at 6,5x10^5^ cells/ml per well in R5 medium and rested overnight at 37°C and 5% CO_2_. Naïve macrophages (M0) were either pre-treated with 500 μg/ml *KKC* extract for 2 hours, then polarized into M1 with 100 ng/ml LPS (Ultrapure; InvivoGen) and 20 ng/mL IFNγ (R&D); M2a with 20 ng/ml IL-4 (R&D) or M2c with 20 ng/ mL IL-10 (R&D) for 22 hours or left unstimulated in accordance with the literature (Iqbal and Kumar, 2015; Purcu et al., 2022; Raggi et al., 2017; Wang et al., 2014).

### 2.5. Flow Cytometry

Single cell suspensions were obtained by detaching the cells from the culture plates using StemPro Accutase Cell Dissociation Reagent (Gibco) according to the manufacturer’s instructions. Exclusion of dead cells and determination of cell viability was performed using Zombie UV Fixable Viability Kit (BioLegend). Surface antigen staining was performed in flow cytometry staining buffer (PBS supplemented with 1% bovine serum albumin, 0.1% sodium azide). Fcγ receptors were blocked prior to surface antigen staining by incubating the cells in flow cytometry staining buffer containing Human TruStain FcX antibody (BioLegend) for 15 minutes on ice. Then, surface staining was performed using following fluorochrome labelled antibodies (Biolegend) in flow cytometry staining buffer at given concentrations for 45 minutes on ice in dark: CD86-BV605 (IT2.2; 1:200), CD38-BV510 (HB-7; 1:100), HLA-DR-APC-Cy7 (L243; 1:200), CD64-BV421 (OX-108, 1:200.), CD206-AF700 (15.2; 1:200), CD209-PE (9E9A8; 1:200), CD200R-PE-Dazzle594 (OX108; 1:200), CD163-BV510 (GHI/61; 1:200). The cells were stained in V-bottom 96-well plates (Greiner Bio-One). Following the staining process, samples were washed, resuspended in flow cytometry buffer and transferred in FACS tubes (Greiner Bio-One). Fluorescence was detected with LSR Fortessa (BD Biosciences). The data were analyzed with FlowJo software (TreeStar).

### 2.6. Total RNA isolation and Real-time PCR

Total RNA isolation of samples was performed using Monarch Total RNA Miniprep Kit (New England Biolabs) according to the manufacturer’s protocol. The purity and quantity of RNA were assessed by Nanodrop spectrophotometer (ThermoFisher Scientific). 1000 ng of RNA per sample was converted to cDNA using EvoScript Universal cDNA Master (Roche) according to the manufacturer’s instructions. M1 and M2 polarization markers, Type I IFN and housekeeping gene expressions were assessed using Fast Start Essential DNA Green Master (Roche) and, RealTime ready Single Assay Primers for IDO1 (#QT00000504), IL1β (QT00021385), IL12A (QT00000357), TNF (QT00029162), TGM2 (QT00081277), MRC1 (CD206) (QT00012810), CD163 (QT00074641), and IL-10 (QT00041685). Real-time PCR analysis was performed using Applied Biosystems 7500 Fast Real-Time PCR System (ThermoFisherScientific). For normalization, housekeeping gene β-actin (ACTB) (QT00095431) was used. 2^-ΔΔCt^ method was used to calculate relative mRNA expression (Livak and Schmittgen, 2001).

### 2.7. ELISA

Supernatants of samples were collected after indicated stimulation times and stored at -20°C until further usage. Concentrations of cytokines were determined using the following kits according to the manufacturer’s instructions: IL-12p70 (Biolegend, 431704), TNF (Biolegend, 430204), and IL-10 (Biolegend, 430604).

### 2.8. Phagocytosis assay

For the assessment of phagocytosis, pHrodo Red conjugated *Staphylococcus aureus* (*S.aureus*) Bioparticle (Invitrogen, A10010) was used. Macrophages were seeded in 24-well cell culture plates at 6,5x10^5^ cells/ml per well in R5 medium. After indicated stimulation times, supernatants were collected, cells were washed with PBS and detached from the culture plate using StemPro Accutase Cell Dissociation Reagent (Gibco) according to the manufacturer’s instructions. After washing, macrophages were incubated with pHrodo Red conjugated *Staphylococcus aureus (S.aureus)* at 37°C (and 4°C for control) for 2 hours. Then, Zombie UV Fixable Viability Kit (BioLegend) staining was performed to eliminate dead cells, and phRodo red fluorescence emission was detected by flow cytometry using LSR Fortessa (BD Biosciences). The data was analyzed with the FlowJo software (TreeStar). Doublets were excluded from the analysis using forward- and side-scatter parameters.

### 2.9. Statistical analyses

Graphpad Prism version 9 (GraphPad) was used to analyze descriptive statistics and to generate graphs. Shapiro-Wilk normality test was used to determine whether the continuous variables are normally distributed. For normally distributed data analyses, two-tailed Student’s t-test was used for two independent samples and one-way ANOVA method was applied for comparison of more than two samples. P values less than 0.05 were considered statistically significant.

## 3. Results

### 3.1. The impact of the KKC extract on the viability of primary human macrophages

First, we analyzed the effect of *KKC* on the viability of primary human macrophages. The cells were either treated with *KKC* extract at concentrations of 50 μg/mL, 100 μg/mL, 500 μg/mL, or 1000 μg/mL for 24 hours or left untreated. After that the cells were stained with Ghost UV and analyzing by flow cytometry. Figure 1 shows that the cell viability was not significantly affected by *KKC* extract at the concentrations of 50 μg/mL, 100 μg/mL, and 500 μg/mL. Although treating the macrophages with 1000 μg/mL *KKC* decreased cell viability significantly compared to untreated control (p=0,0134 = *), more than 78 % of the cells were viable (Figure 1).

**Figure 1.**
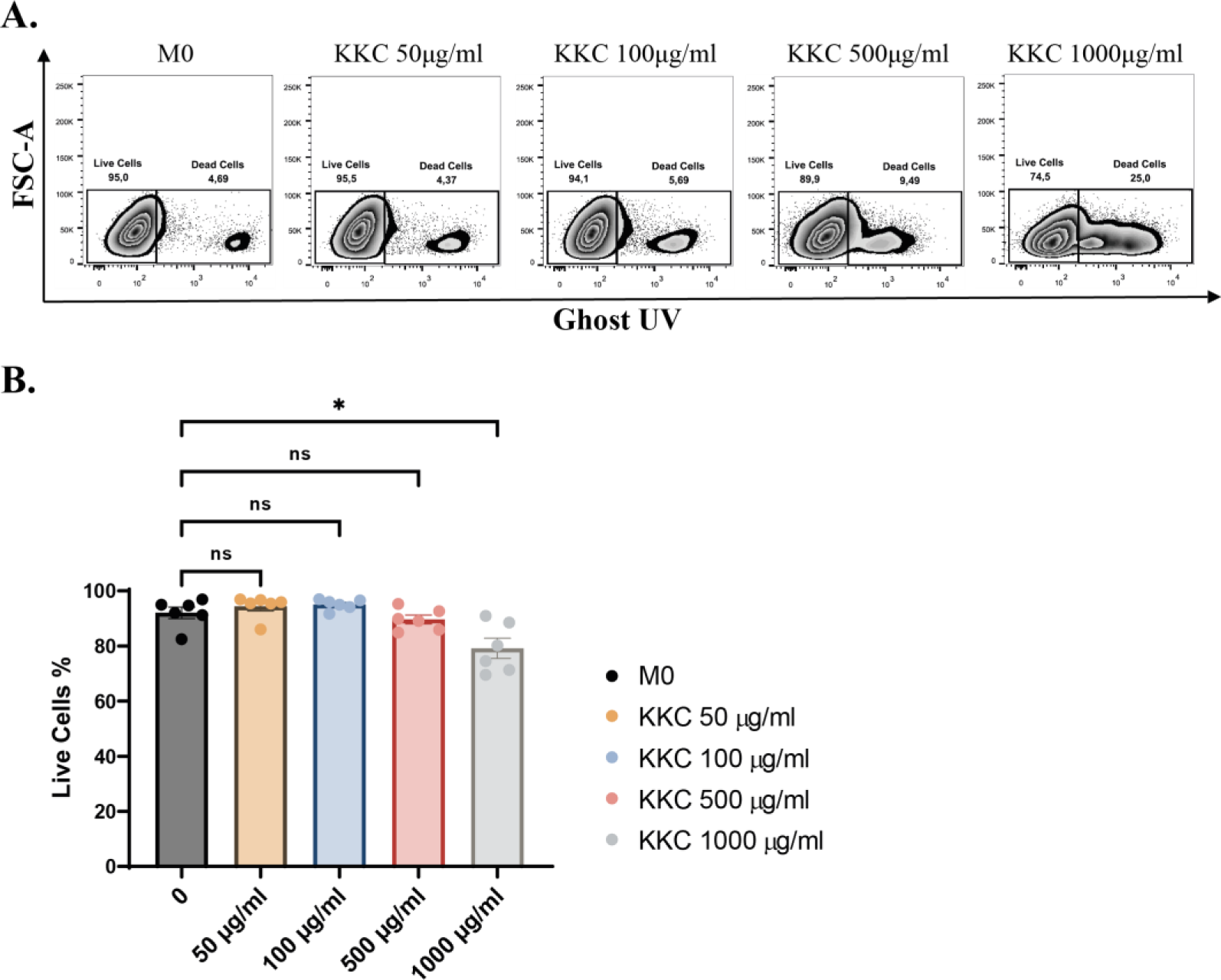
The effect of *KKC* extract on the viability of human MDMs. Unpolarized primary human MDMs (M0) were either treated with the *KKC* extract at doses of 50 μg/mL, 100 μg/mL, 500 μg/mL, or 1.000 μg/mL for 24 h or left untreated. Macrophages were stained with a Zombie UV fixable viability kit (Biolegend) and analyzed by flow cytometry. A) Representative dot blots, B) The bar graphs show the percentages of live cells. Data shown are mean ± SEM of biological replicates of 6 donors pooled from 3 independent experiments. One-Way ANOVA followed by Dunnet’s post-hoc test was performed for the statistical analyses. *P<0.05 / p=0.0134 (*).

### 3.2. The impact of the KKC extract on the expression of M1 and M2 markers in primary human macrophage phenotypes at mRNA level

Next, we examined how the *KKC* extract affects the expression of conventional M1 and M2 macrophage markers, which are widely accepted in the literature (Martinez et al., 2006; Martinez and Gordon, 2014; Nadella et al., 2015; Rőszer, 2015; Tóth et al., 2009). The relative mRNA levels of M1 genes, IL-1β, IL-12a and TNF was increased in M0, M1, M2a and M2c macrophages treated with 500 μg/mL *KKC* extract (Figure 2). IDO1 expression levels were also increased in macrophage subtypes except in M1 macrophages (Figure 2). In contrast, the relative expression levels of M2 genes CD163 and IL-10 were decreased in *KKC* treated M0, M2a and M2c macrophages (Figure 2A-C-D). Intriguingly, in M1 macrophages, while the CD163 expression was decreased, IL-10 expression was increased (Figure 2B). Moreover, the TGM2 expression level was increased in *KKC* treated M0, M1, M2a and M2c macrophages. Although TGM2 is widely accepted as an M2 marker, there are contradictory studies suggesting that its expression may be induced in some inflammatory conditions (Sun and Kaartinen, 2018). Interestingly, after the treatment with the *KKC* extract, while the expression level of another M2 marker MRC-1 was increased in M0 and M1 macrophages (Figure 2A-B), it decreased in M2a and M2c macrophages (Figure 2C-D). Our data show that the *KKC* extract increased the expression of M1 markers in unpolarized and polarized macrophages at mRNA level. However, it had increasing or decreasing effects depending on the M2 markers and the macrophage subtypes.

**Figure 2.**
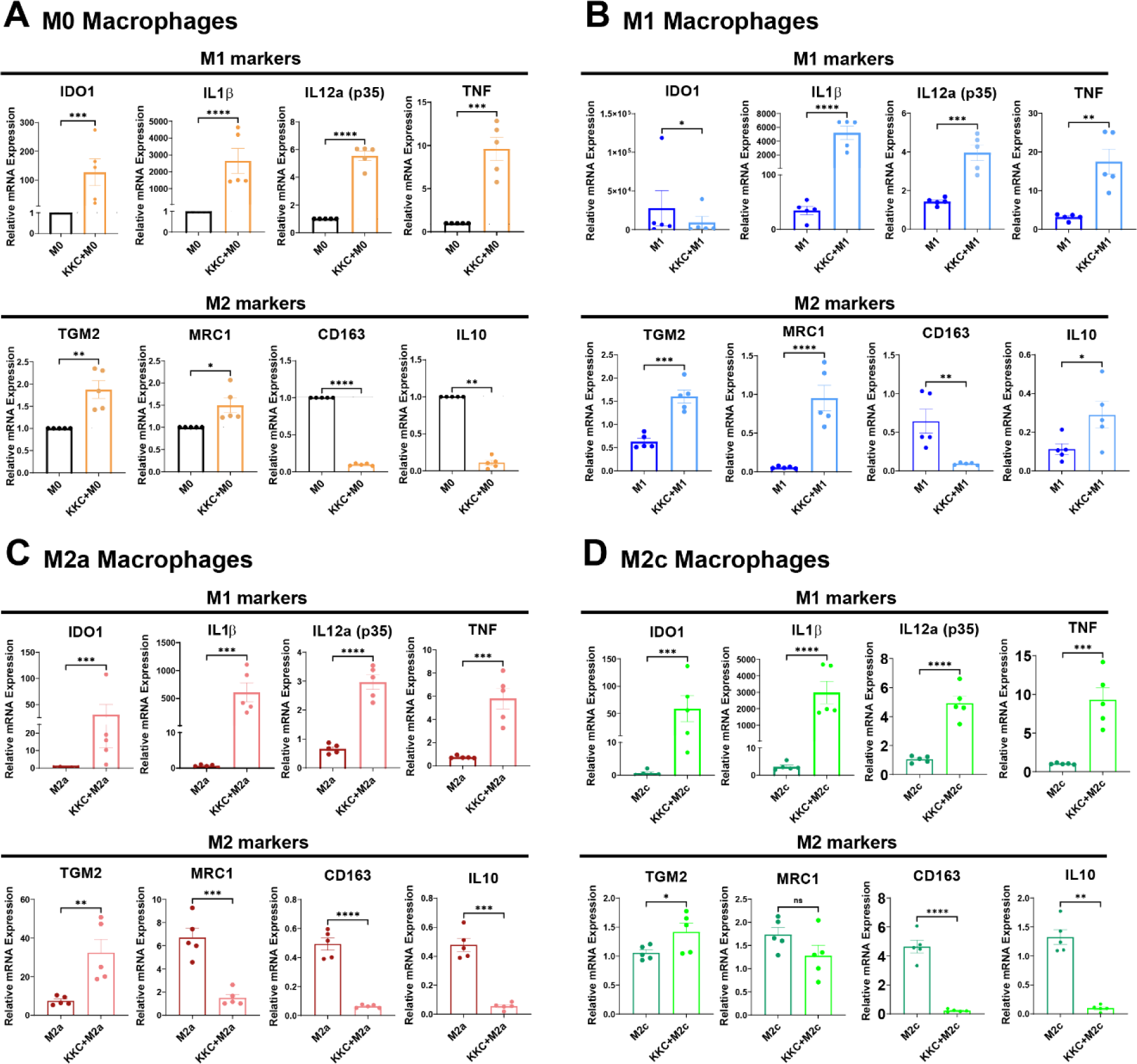
The effect of *KKC* extract on the expression of M1/M2 markers by unpolarized and polarized human MDMs at mRNA level. Unpolarized primary human MDMs (M0) were only treated with 500 μg/mL *KKC* extract for 24 h. For polarization, human MDMs were pre-stimulated with 500 μg/mL *KKC* extract for 2 h and then polarized into M1 (100 ng/ml and 20 ng/ml IFNγ), M2a (20 ng/ml IL-4) and M2c (20 ng/ml IL-10) phenotypes for 22 h. Polarization controls were only stimulated with M1, M2a or M2c agents. qPCR analysis of common M1 and M2 markers in (A) M0, (B) M1, (C) M2a, and (D) M2c macrophages were shown. Data shown are mean ± SEM of biological replicates of 5 donors pooled from 3 independent experiments. Two-tailed paired Student’s t-test was performed for the statistical analyses comparing controls and *KKC* extract treated groups. *P<0.05, **P<0.01, ***P<0.001, ****P<0.0001.

### 3.3. The impact of the KKC extract on the expression of M1 and M2 markers in primary human macrophage phenotypes at protein level

Next, we examined the impact of *KKC* extract on the expression of conventional M1 and M2 macrophage markers (Gordon, 2003; Italiani and Boraschi, 2014; Martinez and Gordon, 2014; Murray et al., 2014) at the protein level. Figure 3 shows the effect of the *KKC* extract stimulation on the expression of M1/M2 surface markers in M0 macrophages. *KKC* treatment did not affect the expression of M1 markers HLA-DR, CD86, CD38; and M2 markers CD200R, CD206, CD209 (Figure 3A-B-D-E-F-G). However, a statistically significant decrease in the expression of the M1 marker CD64 and M2 marker CD163 was observed after *KKC* treatment compared to control (Figure 3C-H).

**Figure 3.**
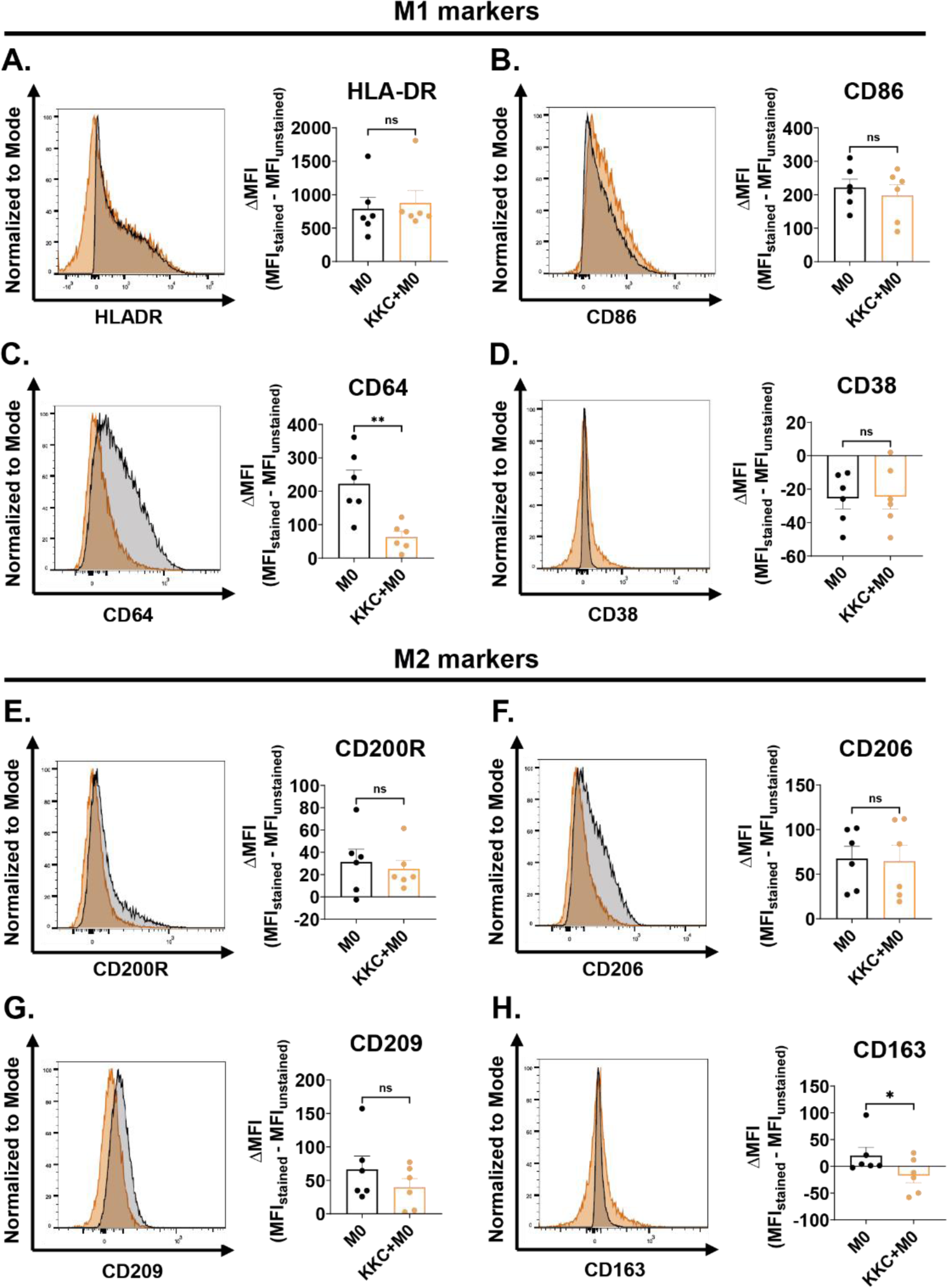
The effect of *KKC* extract stimulation on the expression of M1/M2 surface markers in M0 macrophages. Unpolarized primary human MDMs (M0) were only treated with 500 μg/mL *KKC* extract or treated with 500 μg/mL DMSO as control for 24 h. (A-D) Expression of HLA-DR, CD86, CD64 and CD38 M1 markers, (E-H) CD200R, CD206, CD209 and CD163 M2 markers were analyzed by flow cytometry. Histograms are given as representative. Data shown are mean ± SEM of biological replicates of 6 donors pooled from 3 independent experiments. Two-tailed paired Student’s t-test was performed for the statistical analyses comparing controls and *KKC* extract treated groups. *P<0.05, **P<0.01, ***P<0.001, ****P<0.0001. MFI: median fluorescence intensity.

The impact of *KKC* extract stimulation on M1/M2 surface marker expression in M1 macrophages is demonstrated in Figure 4. M1 macrophages showed a significant decrease in the expression of the M1 markers HLA-DR, CD86, CD64, CD38; and M2 marker CD209 after *KKC* stimulation compared to control. (Figure 4A-B-C-D-G). Nevertheless, the expression levels of M2 markers CD200R, CD206, and CD163 did not change in M1 macrophages after treatment with the *KKC* extract (Figure 4E-F-H).

**Figure 4.**
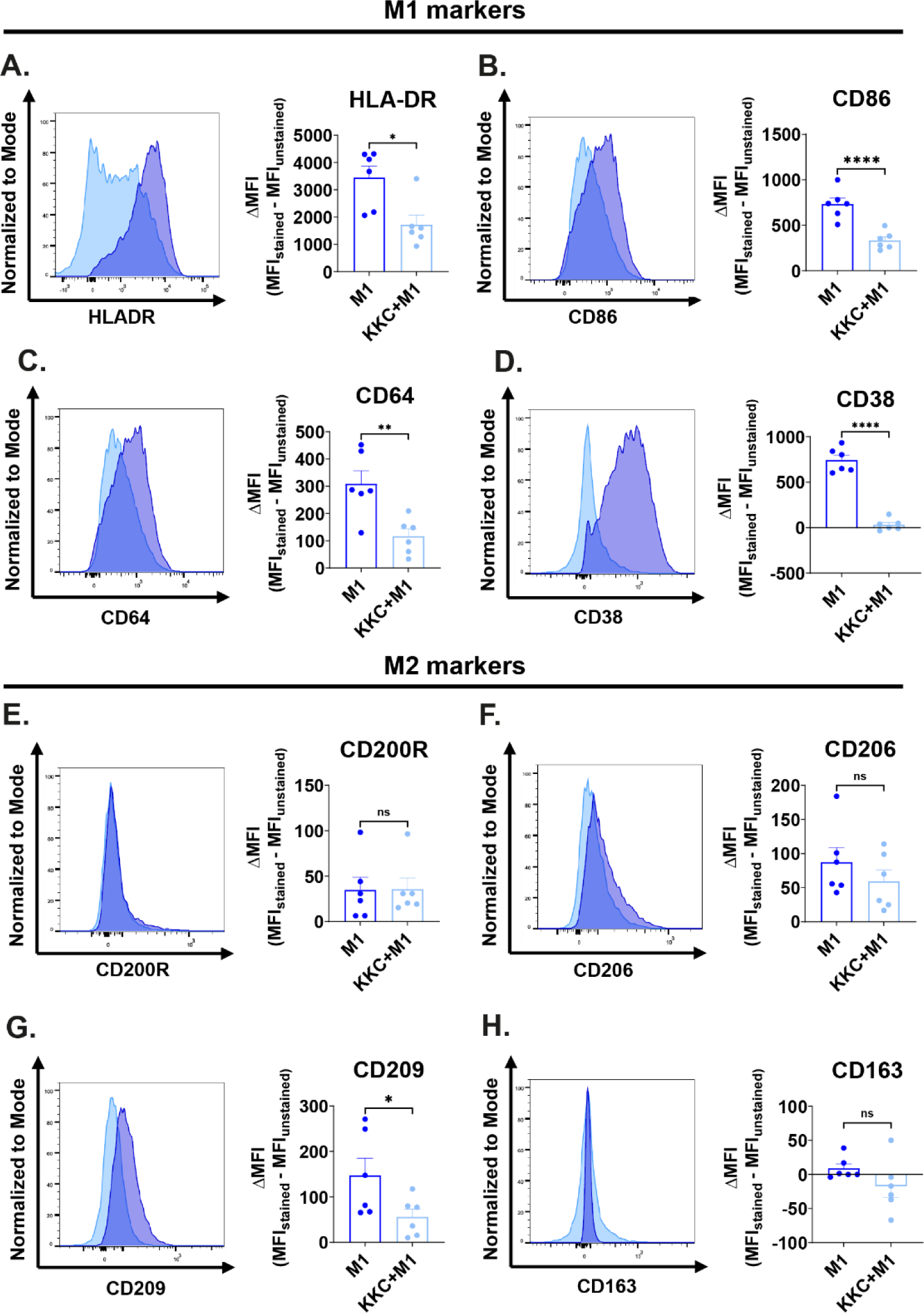
The effect of *KKC* extract stimulation on the expression of M1/M2 surface markers in M1 macrophages. Primary human macrophages were pre-stimulated with 500 μg/mL *KKC* extract for 2 h and then polarized into M1 (100 ng/ml LPS + 20 ng/ml IFNγ) macrophages (*KKC* group macrophages) for 22 hours. Polarization controls were treated with 500 μg/mL DMSO for 2 h and with M1 agents for 22 h. (A-D) Expression of HLA-DR, CD86, CD64, and CD38 M1 markers and (E-H) CD200R, CD206, CD209 and CD163 M2 markers were analyzed by flow cytometry. Histograms are given as representative. Data shown are mean ± SEM of biological replicates of 6 donors pooled from 3 independent experiments. Two-tailed paired Student’s t-test was performed for the statistical analyses comparing controls and *KKC* extract treated groups. *P<0.05, **P<0.01, ***P<0.001, ****P<0.0001. MFI: median fluorescence intensity.

Figure 5 shows the effect of the *KKC* extract stimulation on M1/M2 surface marker expression in M2a macrophages. *KKC* treatment did not significantly affect the expression of the M1 markers HLA-DR, CD86, CD38 and M2 markers CD209, CD163 (Figure 5 A-B-D-G-H). However, a statistically significant decrease in the expression of M1 marker CD64; and M2 markers CD206 and CD200R (Figure 5C-F-E) was observed in *KKC* treated M2a macrophages compared to M2a control macrophages.

**Figure 5.**
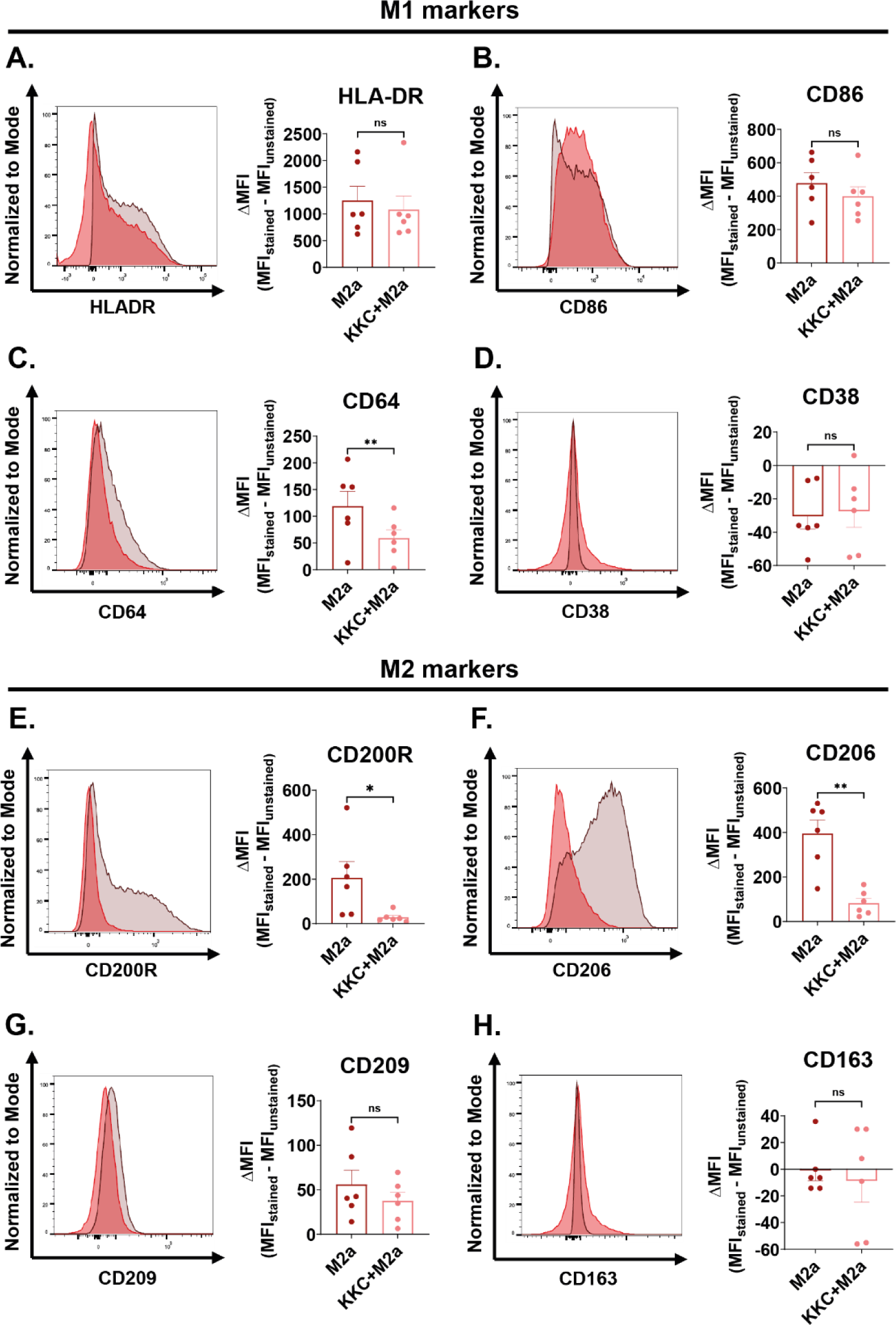
The effect of *KKC* extract stimulation on the expression of M1/M2 surface markers in M2a macrophages. Primary human macrophages were pre-stimulated with 500 μg/mL *KKC* extract for 2 h and then polarized into M2a (20 ng/ml IL-4) macrophages (*KKC* group macrophages) for 22 hours. Polarization controls were treated with 500 μg/mL DMSO for 2 h and with M2a agent for 22 h. (A-D) Expression of HLA-DR, CD86, CD64, and CD38 M1 markers and (E-H) CD200R, CD206, CD209 and CD163 M2 markers were analyzed by flow cytometry. Histograms are given as representative. Data shown are mean ± SEM of biological replicates of 6 donors pooled from 3 independent experiments. Two-tailed paired Student’s t-test was performed for the statistical analyses comparing controls and *KKC* extract treated groups. *P<0.05, **P<0.01, ***P<0.001, ****P<0.0001. MFI: median fluorescence intensity.

Figure 6 shows the effect of the *KKC* extract stimulation on the expression of the M1/M2 surface markers in M2c macrophages. After the *KKC* treatment, a statistically significant reduction in the expression of the M1 markers HLA-DR and CD64; and M2 markers CD209 and CD163 was observed (Figure 6A-C-G-H) in M2c macrophages compared to untreated control. *KKC* treatment did not significantly affect the expression of the M1 markers CD86 and CD38 and M2 markers CD200R and CD206 (Figure 6 B-D-E-F).

**Figure 6.**
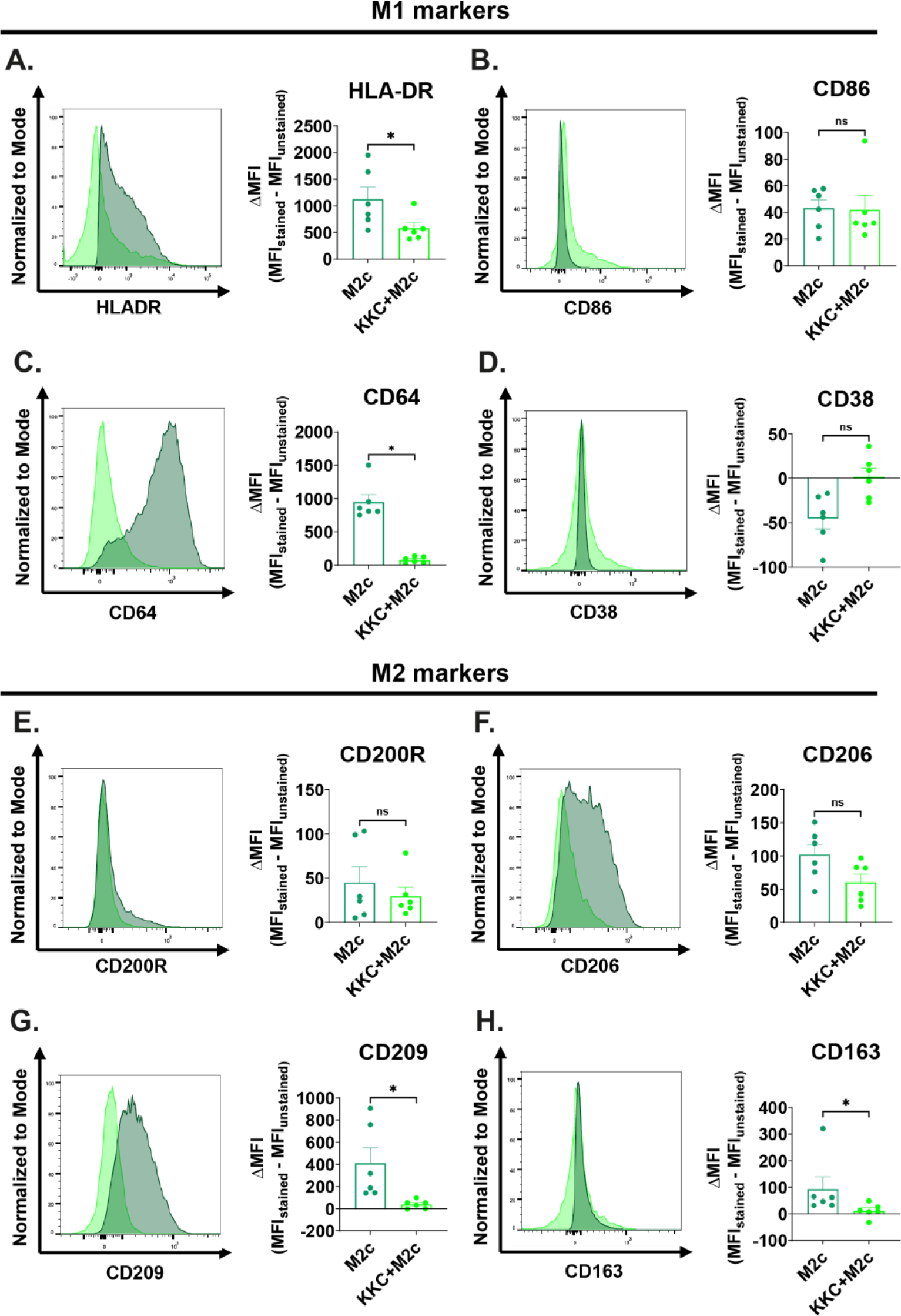
The effect of *KKC* extract stimulation on the expression of M1/M2 surface markers in M2c macrophages. Primary human macrophages were pre-stimulated with 500 μg/mL *KKC* extract for 2 h and then polarized into M2c (20 ng/ml IL-10) macrophages (*KKC* group macrophages) for 22 hours. Polarization controls were treated with 500 μg/mL DMSO for 2 h and with M2c agent for 22 h. (A-D) Expression of HLA-DR, CD86, CD64, and CD38 M1 markers and (E-H) CD200R, CD206, CD209 and CD163 M2 markers were analyzed by flow cytometry. Histograms are given as representative. Data shown are mean ± SEM of biological replicates of 6 donors pooled from 3 independent experiments. Two-tailed paired Student’s t-test was performed for the statistical analyses comparing controls and *KKC* extract treated groups. *P<0.05, **P<0.01, ***P<0.001, ****P<0.0001. MFI: median fluorescence intensity.

Next, we measured the levels of M1 cytokines TNF and IL-12, as well as M2 cytokine IL-10 in M0, M1, M2a, and M2c macrophages by ELISA. As shown in Figure 7, the *KKC* extract treatment increased the production of TNF in M0, M2a, and M2c macrophages, while it did not further increase the production of TNF by M1 macrophages (Figure 7A). There was no IL-12 (p70) production in M0, M2a, and M2c macrophages. However, there was IL-12 (p70) production in M1 macrophages, and its level was decreased after *KKC* extract treatment compared to untreated M1 macrophages (Figure 7B). Interestingly, IL-10 showed a unique pattern of response, a decrease in mRNA expression (Figure 2A-C) but increase in secretion in M0 and M2a macrophages (Figure 7C), while the opposite response was observed in M1 macrophages (Figure 2B-D, 7C).

**Figure 7.**
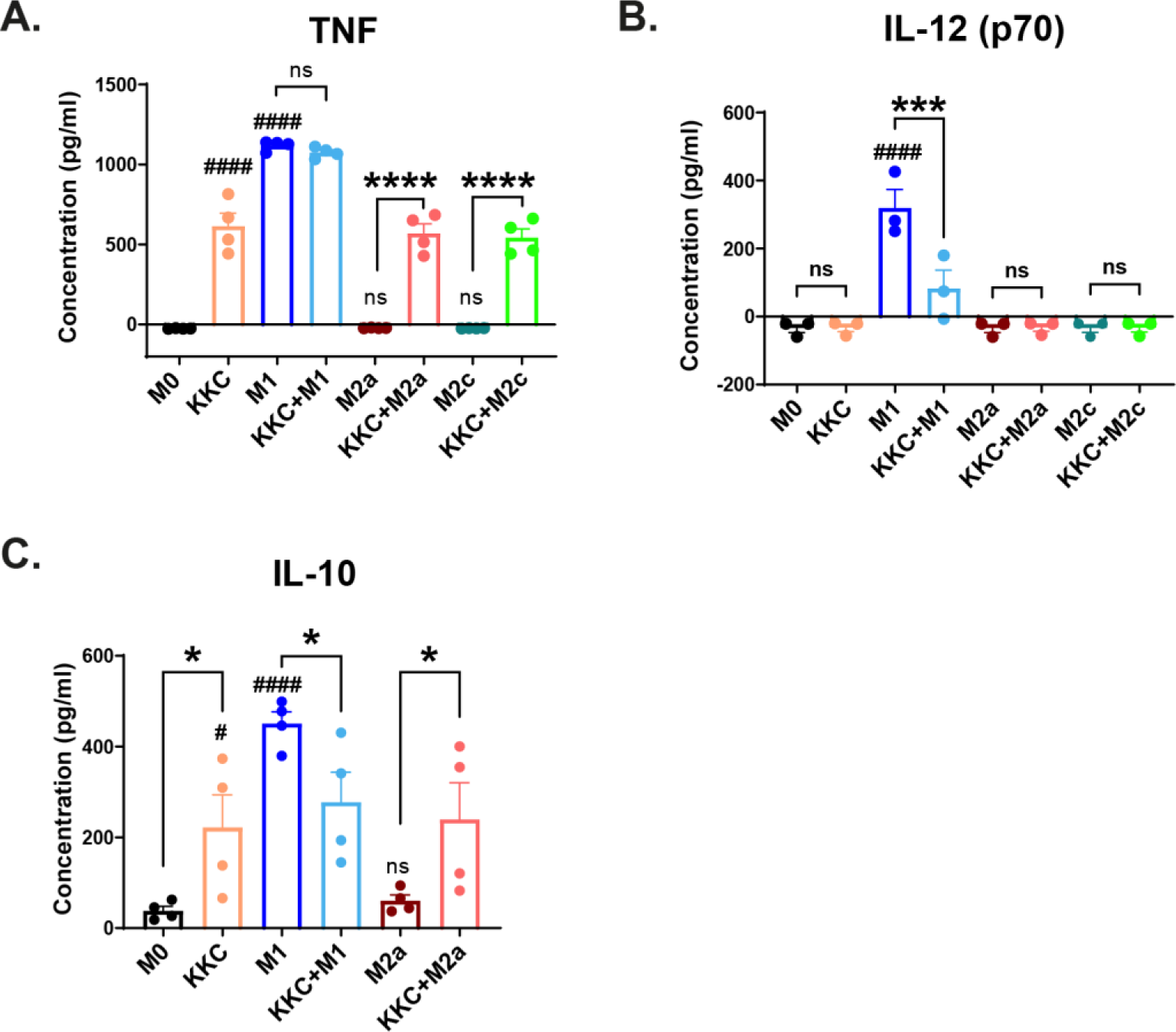
The effect of *KKC* extract on M1/M2 cytokine production by polarized and unpolarized MDMs. Unpolarized primary human MDMs (M0) were only treated with 500 μg/mL *KKC* extract for 24 h. For polarization, human MDMs were pre-stimulated with 500 μg/mL *KKC* extract for 2 h and then polarized into M1 (100 ng/ml and 20 ng/ml IFNγ), M2a (20 ng/ml IL-4) and M2c (20 ng/ml IL-10) for 22 h. Polarization controls were only stimulated with M1, M2a or M2c agents. ELISA analysis of (A) TNF (B) IL-12 (p70) and (C) IL-10 were shown. Data shown are mean ± SEM of biological replicates of 6 donors pooled from 3 independent experiments. One-Way ANOVA followed by Sidak’s post-hoc test was performed for the statistical analyses comparing controls and *KKC* extract treated groups. # indicate p values obtained by comparing M1, M2a and M2c with M0 control. * indicate p values obtained by comparing M1 vs. *KKC*+M1, M2a vs. *KKC*+M2a, and M2c vs. *KKC*+M2c. *, # P<0.05, **, ##P<0.01, ***, ### P<0.001, ****, ####P<0.0001.

Overall, our data suggest that *KKC* extract, while increasing the levels of some of the M1 markers, mostly acts on the modulation of the macrophage inflammatory response.

### 3.4. The impact of the KKC extract on the phagocytosis capacity of primary human macrophage phenotypes

During our investigation into the effect of *KKC* extract on macrophage polarization, we discovered a decrease in the protein expression levels of surface markers CD64, CD206, CD209 and CD163 in the M0, M1, M2a, and M2c macrophage groups (Figure 2-6). While these markers are commonly linked to the polarization of macrophages, they also have a significant impact on the phagocytic activity of these cells (Akinrinmade et al., 2017; Azad et al., 2014; Bournazos et al., 2016; Etzerodt and Moestrup, 2013; Fabriek et al., 2005; Martinez et al., 2008; McGreal et al., 2005). Hence, the effect of the *KKC* extract on the phagocytic capacity of human MDMs was analyzed using pHRodo conjugated heat-killed *Staphylococcus aureus* bioparticle. Our data showed that the *KKC* extract prominently decreased the phagocytic capacity of unpolarized and polarized macrophages (Figure 8).

**Figure 8.**
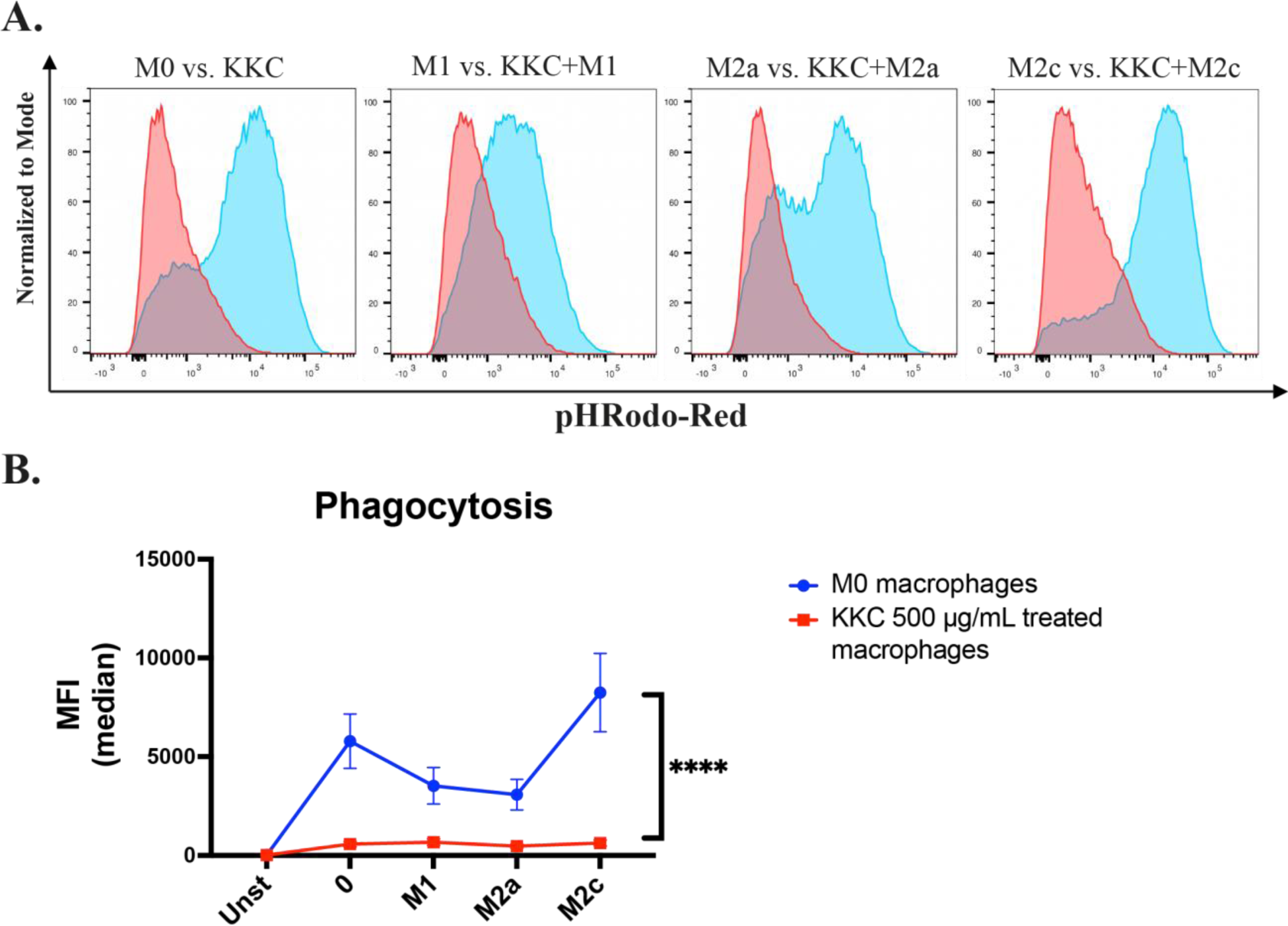
The effect of *KKC* extract on the phagocytic activity of polarized and unpolarized human MDMs. Unpolarized primary human MDMs (M0) were only treated with 500 μg/mL *KKC* extract for 24 h. For polarization, human MDMs were pre-stimulated with 500 μg/mL *KKC* extract for 2 h and then polarized into M1 (100 ng/ml and 20 ng/ml IFNγ), M2a (20 ng/ml IL-4) and M2c (20 ng/ml IL-10) for 22 h. Polarization controls were only stimulated with M1, M2a or M2c agents. Macrophages were incubated with pHrodo Red-tagged *S.aureus* bioparticle for 2 h at 37^0^C. Fluorescence in PE-channel was analyzed by flow cytometry. A) Representative histograms, B) The line graphs show the pHRodo-Red positive MFIs. Data shown are mean ± SEM of biological replicates of 6 donors pooled from 3 independent experiments. Two-way ANOVA was performed for the statistical analyses comparing controls and *KKC* extract treated groups.

## 4. Discussion

In the last 20 years, medicinal plant research has gained much importance. Many studies have shown that plants and their chemicals can prevent and treat a variety of diseases including cancer, and infectious diseases caused by agents like avian and swine influenza virus. The polyherbal formulation, *Kabasura kudineer choornam* (*KKC*), comprised of 15 ingredients (Behera et al., 2023; John et al., 2015; Mekala and Murthy, 2020), have been shown to have anti-inflammatory, anti-pyretic and anti-bacterial activities (John et al., 2015; Mekala and Murthy, 2020; Natarajan et al., 2020; Thillaivanan et al., 2015). Besides, related with the recent SARS-CoV2 outbreak, it has been shown that *KKC* has high binding affinity and interactions with SARS-CoV-2 spike protein shown in *in-silico* studies (Kiran et al., 2020; Kumar et al., 2020; Maideen, 2021; Nallusamy et al., 2021; Vincent et al., 2020) and exhibits good anti-viral properties against SARS-CoV-2 shown in a few clinical studies (Meenakumari et al., 2021; Natarajan et al., 2020; Srivastava et al., 2021, 2021). Following these intriguing and promising results, a limited number of studies were conducted to investigate the effect and mechanism of action of the *KKC* on the immune response. For this reason, Jose and their colleagues have demonstrated that the aqueous extract of the *KKC* has an antioxidant and anti-inflammatory effect by inhibiting the production of intracellular ROS and NO and decreasing the production of pro-inflammatory cytokines in a mouse macrophage cell line (Jose et al., 2022). Furthermore, Behera and their colleagues investigated the effect of the *KKC* on immunomodulation using Jurkat T cells. They have found that the *KKC* increased the expression of anti-inflammatory IL-10Rα and IL-10 cytokine in LPS induced T cells, indicating a potential role in immunomodulation (Behera et al., 2023). To the best of our knowledge, currently there is no study that examines the impact of *KKC* on macrophage polarization.

In the presence of certain polarization factors, macrophages can be polarized into pro-inflammatory M1 macrophages and anti-inflammatory M2 macrophages (Italiani and Boraschi, 2014; Yang et al., 2014). Disrupted balance between these phenotypes can cause progression of diseases such as cancer, infectious and inflammatory diseases (Espinoza-Jiménez et al., 2012; Kraakman et al., 2014; Menghini et al., 2012; Murray et al., 2014). M1 macrophages have been linked to a strong type I interferon response and elevated disease severity and mortality in infections caused by viruses like SARS-CoV-2 (Feng et al., 2020; Lv et al., 2021; Mehta et al., 2020; Merad and Martin, 2020; Ziegler et al., 2020) and the highly virulent H5N1 influenza A virus (Baskin et al., 2009; Channappanavar et al., 2016). In such infections, the severity of the disease is determined by the severity of the “cytokine storm,” which is caused by the overproduction of pro-inflammatory mediators and the overstimulation of the inflammatory response. Macrophages are considered to play a significant role in this process (Jafarzadeh et al., 2020; Kosyreva et al., 2021). Achieving a balanced activation of the M2-like phenotype is crucial to restrict the immunopathological response caused by the infection (Sang et al., 2015; Shirey et al., 2014). Therefore, identifying modulators regulating macrophage polarization is crucial for effective immunotherapies. Effect of *KKC* on macrophage polarization is not known. Here, we report a detailed analysis of the impact of *KKC* on the polarization and function of primary human monocyte-derived macrophages.

To determine the effect of the *KKC* extract on human macrophage polarization, first we analyzed the conventional polarization markers at mRNA level. We found that although the expression of the M1 markers was increased in unpolarized M0, M1, M2a and M2c polarized macrophages, the expression of TGM2 which is an M2 marker was also increased (Figure 2). Although TGM2 is widely accepted as an M2 marker (Martinez et al., 2013), there are contradictory studies suggesting that TGM2 expression may be induced in some inflammatory conditions (Kim et al., 2006; Sun and Kaartinen, 2018; Verma and Mehta, 2007). TGM2 also could activate TGF-β, which in turn induces profibrotic and anti-inflammatory cytokines (Mehta et al., 2010). More importantly, TGM2 plays a crucial function in efferocytosis (Eligini et al., 2016), a process essential for tissue repair, inflammation resolution, and maintaining immune system balance in homeostasis (Ge et al., 2022). Thus, it can be inferred that the induction of TGM2 contributes to the immunomodulatory effect of *KKC* (Figure 2). Moreover, IL-10 expression was found to be decreasing in unpolarized M0 and M2a and M2c polarized macrophages; however, increasing in M1 polarized macrophages at mRNA level (Figure 2). This result was in part aligned with the literature (Behera et al., 2023).

Interestingly, an opposite effect was observed at protein level. The secretion of IL-10 was increased in M0 and M2a macrophages, while it was decreased in M1 macrophages after the *KKC* treatment (Figure 7). Hence, the regulation of IL-10 expression could also be considered as a part of the immunomodulatory effect of the *KKC*.

Macrophages are professional phagocytes (Aderem and Underhill, 1999). They express variety of receptors to facilitate this function (Hirayama et al., 2017). M1 and M2 macrophages also differ in their phagocytic activity (Leidi et al., 2009). M2 macrophages have higher phagocytic capacity than M1 macrophages (Gratchev et al., 2005; Lingnau et al., 2007). However, this is highly dependent on the expressed surface markers such as CD206, CD163 and CD209 (Schulz et al., 2019) and the environmental conditions. For instance, the Fc gamma receptor (FcγRI), CD64, has been reported as an M1 polarization marker in the literature (Murray et al., 2014) while having a fundamental role in receptor-mediated phagocytosis. Another M1 marker CD38, found to be selectively upregulated after IFNγ treatment (Schulz et al., 2019), also associated with phagocytosis capacity probably due to its function in releasing Ca2+ ions, which could enhance the process of particle engulfment in a manner comparable to Fcγ receptor-mediated phagocytosis (Kang et al., 2012). The CD206 marker is recognized as an M2a polarization marker in the literature (Martinez et al., 2008). This mannose scavenger receptor is also involved in phagocytosis (Azad et al., 2014; Schulz et al., 2019; Xu et al., 2019) CD163 has been identified as an M2c marker (Mantovani et al., 2013; Shapouri-Moghaddam et al., 2018; Wang et al., 2020) and its expression also renders macrophage phagocytosis competent (Fabriek et al., 2005; Schulz et al., 2019). CD209 (DC-SIGN) is a C-type lectin receptor that recognizes high mannose-type N-glycans, binds with high affinity (Geurtsen et al., 2010) and is involved in phagocytosis (McGreal et al., 2005). Schulz and colleagues have discovered that CD209 marker is directly associated with the amount of particle uptake during phagocytosis (Schulz et al., 2019). Importantly, studies have shown that CD209 can directly bind to the S glycoprotein of SARS-CoV-2 and cooperate with ACE2 for viral entry of SARS-CoV via endocytosis (Han et al., 2007; Xu et al., 2020; Yang et al., 2004). It has also been shown that high expression of CD209 triggers an immune response and can lead to a severe response to SARS-CoV-2 infection in cancer patients (Li et al., 2022). When the effect of *KKC* extract on M1/M2 macrophage polarization in primary human macrophages was examined at the protein level, a decrease in the expression levels of phagocytic surface markers CD64, CD206, CD209 and CD163 was observed in M0, M1, M2a and M2c macrophage groups (Figure 3-6). Downregulation of another phagocytic surface marker CD38 was also observed in M1 macrophages. (Figure 3).

The changes observed in the surface marker expressions lead us to investigate the phagocytosis function of polarized and unpolarized macrophages after *KKC* stimulation. As shown in the Figure 8, overall phagocytosis capacity of human monocyte derived macrophages was prominently decreased after the *KKC* treatment, as expected. In infections caused by viruses such as SARS, SARS-CoV-2 (Feng et al., 2020; Lv et al., 2021; Mehta et al., 2020; Merad and Martin, 2020; Ziegler et al., 2020) and the highly virulent H5N1 influenza A virus (IAV) (Baskin et al., 2009; Channappanavar et al., 2016), M1 macrophages have been associated with a high type I interferon response and cytokine storm leading to increased disease severity and mortality (Jamilloux et al., 2020). CD169+ tissue macrophages from patients who died from SARS-CoV-2 infection have been shown to express the SARS-CoV-2 entry receptor ACE2 and contain SARS-CoV-2 nucleoprotein (NP) (Feng et al., 2020; Jalloh et al., 2022). Although, early studies showed that resident macrophages in human tissues do not express ACE2 (Sungnak et al., 2020; Zhao et al., 2020), recently it has been found that a specific subset of peripheral CD14+ monocytes and macrophages of healthy individuals express ACE2 (Zhang et al., 2021). In addition, C-type lectin receptors DC-SIGN (CD209), L-SIGN and antibody-mediated internalization can provide further routes for viral entry of SARS-CoV and SARS-Cov-2 via endocytosis (Amraei et al., 2021; Han et al., 2007; Xu et al., 2020; Yang et al., 2004). Therefore, the decrease in the phagocytosis and phagocytic markers such as CD209, may help to alleviate the disease symptoms. In the light of these information combining with our findings suggest that *KKC* may be a potential immunomodulatory agent against viral infections.

## 5. Conclusions

We report here the impact of the medicinal polyherbal formulation *KKC* extract on polarization and phagocytic activity of primary human macrophages. Altogether, our data suggest that *KKC* extract modulates macrophage inflammatory response and could be a potential supplement for the treatment of infectious and inflammatory diseases. To the extent of our knowledge, this is the first study investigating the effect of the *KKC* extract on primary human macrophage polarization and our data provide useful information on the immunomodulatory abilities of the *KKC* extracts.

## CRediT authorship contribution statement

**Asli Korkmaz:** Formal analysis, Investigation, Conceptualization, Methodology, Validation, Writing—original draft Preparation, Visualization. **Duygu Unuvar:** Methodology, Investigation. **Sinem Gunalp:** Methodology, Investigation. **Derya Goksu Helvaci:** Methodology, Investigation. **Duygu Sag:** Conceptualization, Methodology, Validation, Investigation, Writing—original draft, Visualization, Supervision, Project administration. All authors have read and agreed to the published version of the manuscript.

## Institutional review board statement

The study was conducted in accordance with the Declaration of Helsinki and approved by the Ethics Committee of Izmir Biomedicine and Genome Center, Izmir, Turkiye (protocol code 2021-042).

## Acknowledgments

We thank Dr. Ravi Kumar Reddy of Sri Sri Tattva (Bengaluru, Karnataka, India) for generously supplying the *Kabasura kudineer* powder. We thank Prof. Fahri Saatcioglu for his valuable support. We thank Dr. Yavuz Dogan and the laboratory staff at Dokuz Eylul University Blood Bank for supplying buffy coats. Additionally, we thank staff of the Flow Cytometry Facility at iBG for their technical assistance.

